# Adaptively introgressed Neandertal haplotype at the OAS locus functionally impacts innate immune responses in humans

**DOI:** 10.1101/051466

**Authors:** Aaron J. Sams, Anne Dumaine, Yohann Nédélec, Vania Yotova, Carolina Alfieri, Jerome E. Tanner, Philipp W. Messer, Luis B. Barreiro

## Abstract

The 2’-5’ oligoadenylate synthetase (OAS) locus encodes for three OAS enzymes (OAS1-3) involved in innate immune response. This region harbors high amounts of Neandertal ancestry in non-African populations; yet, strong evidence of positive selection in the OAS region is still lacking. Here we used a broad array of selection tests in concert with neutral coalescent simulations to firmly demonstrate a signal of adaptive introgression at the OAS locus. Furthermore, we characterized the functional consequences of the Neandertal haplotype in the transcriptional regulation of OAS genes at baseline and infected conditions. We found that cells from people with the Neandertal-like haplotype express lower levels of *OAS3* upon infection, as well as distinct isoforms of *OAS1* and *OAS2.* Notably, the Neandertal-introgressed haplotype reintroduced an ancestral splice variant of *OAS1* encoding a more active protein, suggesting that adaptive introgression occurred as a means to resurrect adaptive variation that had been lost outside Africa.

## Background

Whole genome sequencing of several archaic human genomes [1–5] representing Neandertals and an as yet geographically and paleontologically unknown population referred to as Denisovans has revealed gene flow between these populations and the ancestors of present-day humans. Neandertal ancestry makes up approximately 0.5-2 percent of the ancestry of most living humans, with higher amounts of Neandertal ancestry found outside of Africa [6–8]. While it seems that there may have been widespread purifying selection against Neandertal ancestry in humans [6,8,9], some positive selection on Neandertal genes (adaptive introgression) has also been observed [10,11]. Neandertals and other archaic populations inhabited Eurasia for several hundred thousand years [12] and were likely well adapted to their environments. Therefore, some genetic variation inherited from these archaic humans may have been adaptive in modern humans, particularly across phenotypes that are strongly influenced by direct interactions with the surrounding environment [10], such as our immune response to infectious agents [13].

The OAS locus on chromosome 12, which harbors three genes (*OAS1, OAS2, OAS3*) encoding the 2’-5’ oligoadenylate synthetase enzymes has received considerable attention due to its clear signatures of multiple archaic haplotypes in populations outside of Africa [14,15], and the critical role of OAS genes in the innate immune response to viruses [16]. Mendez and colleagues [17] first identified an introgressed haplotype of *OAS1* from Denisovans that is restricted to individuals in Indonesia and Melanesia. Later, these authors [15] identified a Neandertal haplotype at the OAS locus that spans a ~190 kilobase region between two surrounding recombination hotspots.

The elevated frequency of the Neandertal derived alleles in the OAS locus (Figure 1) relative to average levels of Neandertal ancestry in Europeans [6], along with the key role OAS genes play in protective immunity against viral infections raises the possibility that introgressed Neandertal haplotypes at OAS may have been adaptive in modern humans. While some studies provide suggestive evidence of adaptive introgression at the OAS locus [6,11], strong evidence of positive selection in the OAS region is still lacking. Indeed, several studies failed to reject a model of neutral evolution for the Neandertal haplotype when using standard neutrality tests [15,18].

**Figure 1.**
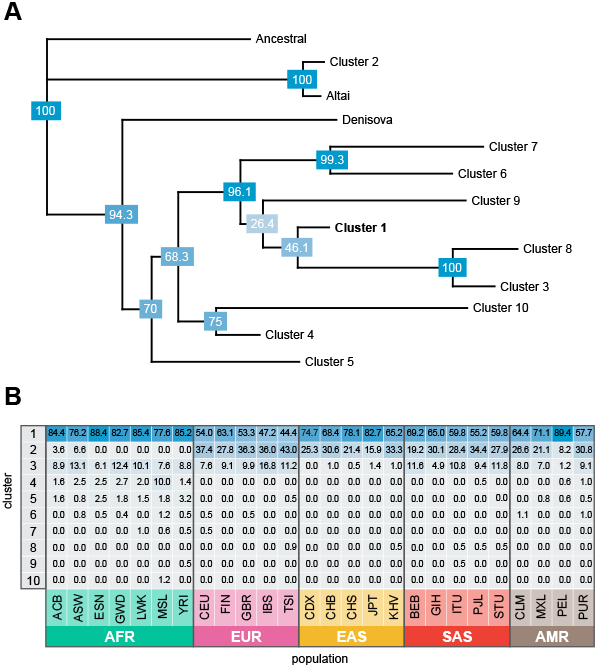
Neandertal introgresssed haplotypes in the OAS region. (A) Neighborjoining tree of 5008 phased haplotypes spanning chr12: 113344739-113449528 (hg19) from phase 3 of the 1000 Genomes Project. Haplotypes were condensed into 10 core haplotypes based on the majority allele after clustering into groups of haplotypes with pairwise differences of 60 or less. The figure illustrates that the Altai haplotype is very similar to “cluster 2” haplotypes found in several human populations, while the Denisovan haplotype is not more closely related to any of the remaining (non-cluster 2) sequences in this dataset. Bootstrap values (1000 replicates) are provided in blue boxes at each node. (B) Frequencies of the 10 core haplotypes within each 1000 Genomes Project population sample. The most common Neandertal-like haplotype, cluster 2, is found only outside of sub-Saharan African samples, with the exception of recently admixed populations. Population codes can be found in Table S7.

We hypothesize that the overall lack of signals of selection in the OAS region stems from the low power of standard neutrality tests to detect adaptive introgression [19]. Here we circumvent this issue by testing the hypothesis of adaptive introgression using extensive neutral coalescent simulations specifically tailored to match the genomic features of the OAS region in combination with several empirical observations of Eurasian genetic variation, and ancient DNA data from Eurasia. We firmly demonstrate a population genetic signal of adaptive introgression at the OAS locus and characterize the functional consequences of the Neandertal haplotype in the transcriptional regulation of OAS genes in macrophages and peripheral blood mononuclear cells (PBMCs) at baseline and infected conditions. We note that the term ‘adaptive introgression’ will be used in a broad sense, as our tests do not allow us to determine the exact timing when selection started to act on the Neanderthal alleles.

## Results

Mendez *et al.* [15] previously reported evidence of archaic introgression at the OAS locus. We first verified these results, using data from phase 3 of the 1000 Genomes Project [20], by examining the relationships of all modern human sequences at the OAS locus with the Altai Neandertal, Denisovan, and an inferred ancestral sequence. Clustering the human haplotypes resulted in 10 consensus sequences representing human haplotype clusters, which we combined with the archaic sequences in a neighbor-joining tree (Methods, Figure 1A, S1). This tree confirms that the Denisovan haplotype is not present in the population samples represented in the 1000 Genomes Project dataset. In contrast, the Neandertal haplotype is found at relatively high frequencies outside of Africa, reaching highest frequencies in European population samples (up to 43%, Figure 1B). Additionally, we corroborate the finding of Mendez and colleagues [15] that Neandertal-like haplotypes in the OAS region are too long to have resulted from incomplete lineage sorting (ILS) (P≤ 2×10^−3^, Methods).

To investigate the hypothesis of non-neutral evolution at the OAS locus we first tested whether the observed frequencies of Neandertal-like sites (NLS) in the OAS region are higher than expected under neutrality. We defined NLS as bi-allelic SNPs with derived alleles that are shared between Neandertals and a non-African population sample, but absent in a sub-Saharan African sample [6,21,22]. Specifically, we simulated the expected allele frequency of NLS under neutrality, using a demographic model based on previously inferred parameters of human demographic history [23–25](Figure S1, Table S1). We considered both a model with a single pulse of Neandertal introgression occurring over a span of 500 years into the ancestral Eurasian population after their population split from Africa, and two additional two-pulse models (Figure S1, Table S1). We found that in all European populations (with the exception of Finnish (FIN)), the highest frequency NLS fall in the extreme 1% of all simulations and are significantly elevated in frequency (all five European samples pass FDR < 0.05, see Table S2), regardless of whether we assume a single-pulse introgression model (Figure 2A) or two-pulse introgression models (Figure S3), indicating that the high frequencies of NLS found in most European populations likely reflects a history of adaptation.

**Figure 2.**
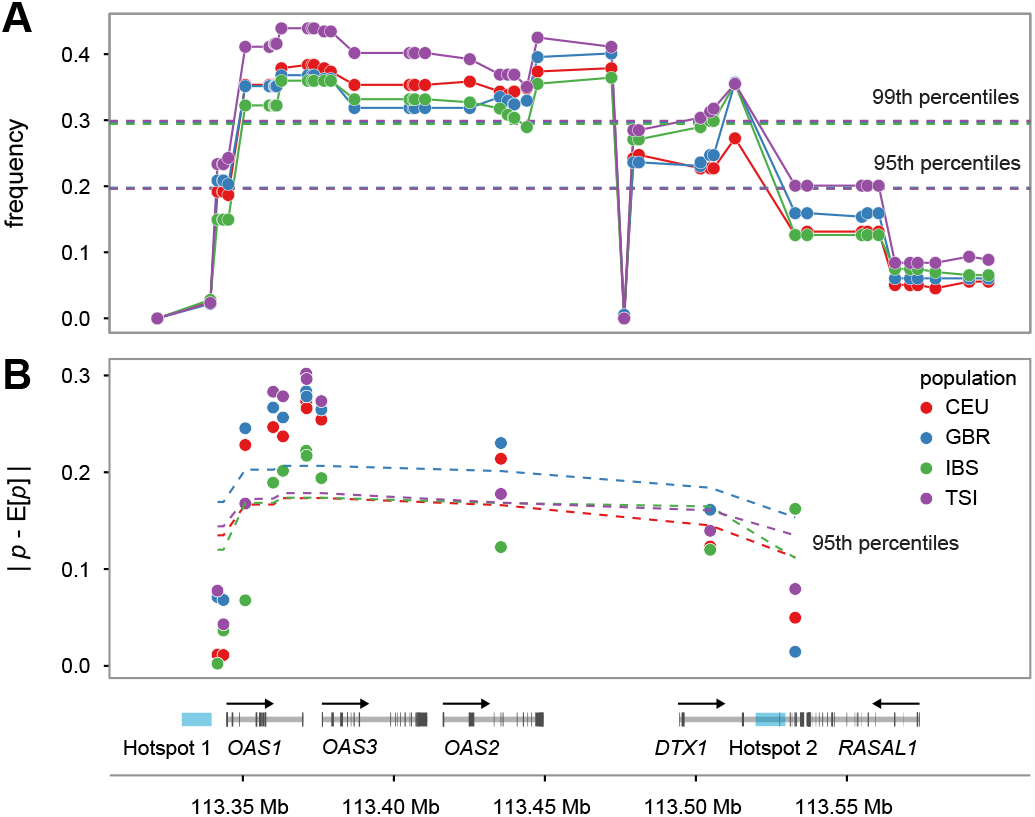
OAS-introgressed haplotypes are found at higher frequencies in European populations than expected under neutrality. (A) Comparison of frequency (y-axis) of NLS in the OAS locus in the CEU, GBR, IBS, and TSI European population samples with respect to neutral expectations (dashed lines) based on coalescent simulations. (B) Absolute difference between observed and expected allele frequency in the same four present-day European samples (y-axis) based on ancient DNA data. Dashed lines represent the 95^th^ percentile of the expected distribution based on similar deviations calculated on a dataset of approximately one million SNPs scattered around the genome and with comparable present-day frequencies to those found for NLS in the OAS region.

We next sought to examine the consistency of the simulation results described above using genetic data from a dataset of 230 ancient Eurasian individuals [26]. Assuming neutrality, the expected frequency of an allele in contemporary European populations can be predicted as a linear combination of allele frequencies sampled from representative ancient populations that have contributed ancestry to present-day European populations in different proportions [26,27] (Methods). Using this approach, we calculated the expected allele frequency in four present day European samples from the 1,000 Genomes Project [20] at the 11 NLS falling within the bounds of the three OAS genes, based on the ancient allele frequencies estimated by Mathieson and colleagues. To set up our null expectations we performed a similar analysis on a dataset of approximately one million SNPs scattered around the genome, generated by Mathieson and colleagues (by merging 213 ancient samples dated between 6,500 and 300 BCE with sequencing data from four European samples from the 1,000 Genomes Project). We found that the NLS at OAS are outliers in the genome with respect to deviations from ancient frequencies. More specifically, we found that the allele frequencies of 6 out of the 11 OAS SNPs tested in the *OAS1-OAS3* region (those 6 SNPs fall within the block of SNPs with NLS frequency greater than 99% of neutral simulations) have increased above the frequency predicted by ancient Eurasian samples by more than 20%, significantly more than what we observed for other SNPs genome-wide with comparable present-day frequencies (lowest *P* = 0.00476, Figure 2B, Table S3). Our findings at this single locus are consistent with results from the genome-wide selection scan performed by Mathieson and colleagues [26] where the OAS region also showed evidence of selection (P < 10^−7^, Table S3), even if it did not reach genome-wide significance after multiple-test correction.

We next searched for additional evidence of recent selection, as measured by the iHS [28] and DIND [29] statistics. These tests share a similar rationale: an allele that has recently been driven to high population frequency by positive selection should be associated with unusually long-range LD (iHS) and reduced intra-allelic nucleotide diversity (DIND) [28–30]. We found that several NLS in the OAS region show significantly high iHS and DIND values with respect to genome-wide expectations (Figure 3A and S4), further supporting that they have been targeted by positive selection.

**Figure 3.**
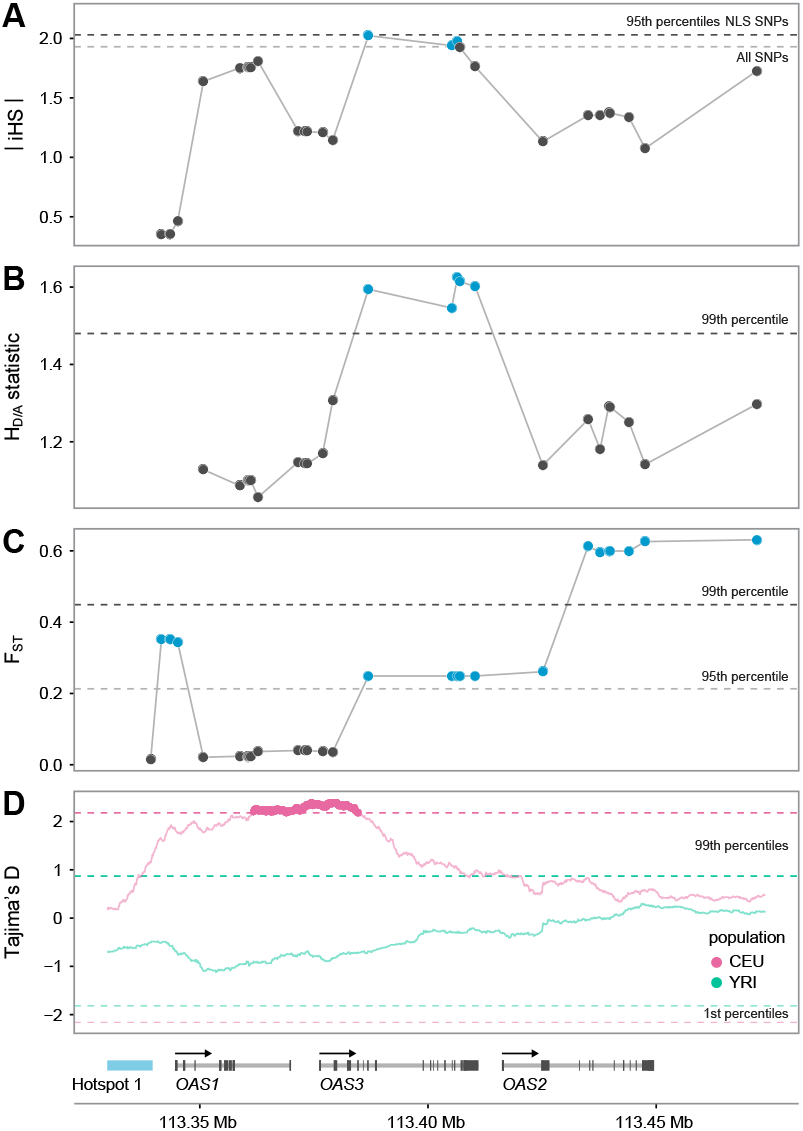
OAS-introgressed haplotypes show multiple signatures of positive selection. (A) Normalized absolute iHS scores (y-axis) across SNPs in the CEU sample. Dashed lines represent the 95^th^ percentile of iHS calculated genome-wide across all SNPs (grey) and all NLS SNPs (black). (B) The *H*_*D/A*_ statistic in CEU (y-axis). Dashed line indicates the 99^th^ percentile of *H*_*D/A*_ calculated across 1,000 simulations. (C) Fst calculated between CEU and all East Asian populations from the 1000 Genomes projects (y-axis). Dashed lines indicate the 95^th^ (grey) and 99^th^ (black) genome-wide percentiles of Fst. Similar results are obtained when comparing any other European population against East Asians (Table S5). (D) Tajima’s D (y-axis) calculated for CEU and YRI samples. Dashed lines indicate the 1^st^ and 99^th^ genome-wide percentiles of Tajima’s D.

As an additional means of understanding whether or not Neandertal haplotypes at OAS are longer than would be expected if they had evolved neutrally, we followed up these empirical haplotype-based tests with a simulation approach using a simple test statistic, *H*_*D/A*_. This statistic (fully defined in Methods) compares pairwise haplotype homozygosity lengths within the set of haplotypes carrying a derived (Neandertal) allele and within those carrying the ancestral allele. We utilized the same demographic models described above for our frequency-based test to understand whether the haplotypes surrounding derived alleles at NLS are longer than expected under a neutral model, conditional on the map of recombination in the OAS region. Again, we found that several derived Neandertal-like alleles are significantly longer than observed in simulated loci, and that these results were robust to a series of alternative simulation models tested, including the one- and two-pulse models described above and two additional one-pulse models which varied the mutation and recombination rates (Figure 3B, Table S4).

Finally, we calculated the levels of population differentiation between European and East Asian populations for all NLS in the OAS region. Interestingly, we found extreme levels of differentiation for most NLS (Fst as high as 0.6, *P*_empirical_ < 0.01, Figure 3C. Table S5), except within the genomic region covering *OAS1* and part of *OAS3*. These results suggest that NLS surrounding *OAS1* have been positively selected in both European and East Asian populations but that distinct haplotypes have been selected in Europe and Asia.

Our population genetic results provide evidence that Neandertal alleles at the OAS locus have likely experienced positive selection during one or several phases after their introduction into the human population, suggesting a possible functional role of these alleles in human innate immune responses. To study this possibility, we analyzed RNA-sequencing data collected on primary macrophages from 96 European-descent individuals, before and after *in-vitro* infection with *Salmonella typhimurium.* After 2 hours of infection, we found that all OAS genes were strongly up-regulated (up to 19-fold, *P* < 1 × 10^−10^, Figure S5), confirming the ability of *Salmonella* to activate the interferon (IFN) production pathway [31–33]. Using genotype data available for the same individuals (673 SNPs spanning the OAS region, see methods) we tested if NLS were associated with variation in the expression levels of *OAS1*, *OAS2* or *OAS3,* in either infected or non-infected macrophages. We found that among NLS, individuals that are heterozygous or homozygous for the Neandertal allele show reduced expression levels of OAS3 (i.e., they were expression quantitative trait loci, or *cis* eQTL for *OAS3*) (Figure 4A, false discovery rate (FDR) < 10%, Table S6). Interestingly, these *cis* eQTL showed a much stronger effect in infected macrophages (best *P*_*salmonella*_ = 3.5×10^−3^ *vs* best *P*_non-infected_ = 0.027), supporting an interaction between the Neandertal haplotype and the *OAS3* response to *Salmonella* infection.

**Figure 4.**
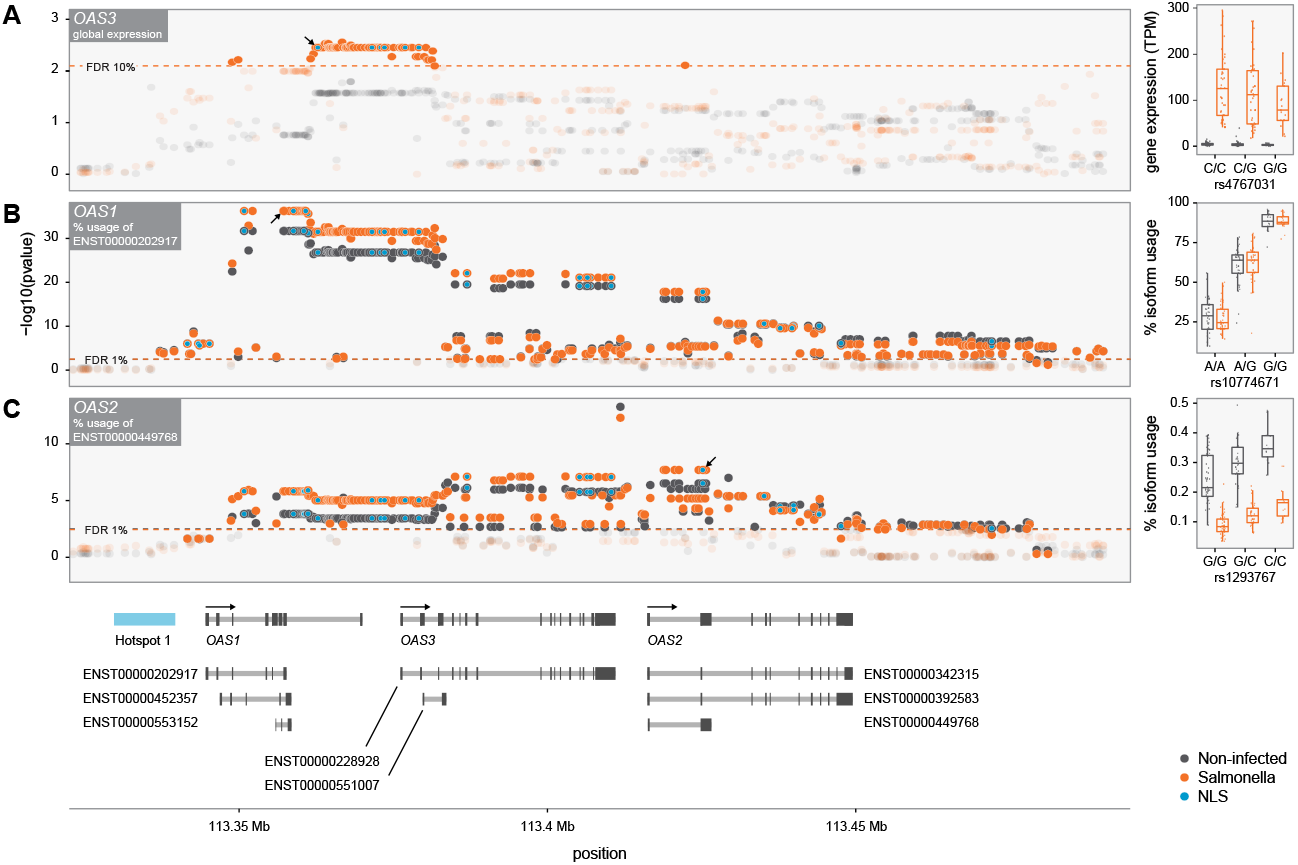
Pervasive impact of the Neandertal haplotype on the regulation of *OAS* genes in primary macrophages. (A) -log 10 Ps (y-axis) for the association between genotypes for SNPs with a MAF > 10% in the OAS region and expression levels of *OAS3* in non-infected (black) and Salmonella-infected macrophages (orange). The dashed line shows the P cutoff corresponding to an FDR of 10%. The right panel shows a boxplot for the association between genotypes at the NLS rs4767031 (x-axis) and the expression levels of *OAS3* in TPM (Transcripts Per Kilobase Million) (y-axis). The lower expression level of *OAS3* in individuals harboring the Neandertal haplotype were confirmed by real-time PCR (Figure S11) (B) -log 10 Ps (y-axis) for the association between genotypes in the OAS regions and the percentage usage of isoform ENST00000202917 (i.e., p46 in the text) in non-infected (black) and Salmonella-infected macrophages (orange). The dashed line shows the P cutoff corresponding to an FDR of 1%. The right panel shows a boxplot for the association between genotypes at the splicing variant rs10774671 (x-axis) and the percentage usage of isoform p46 (y-axis). (C) Similar to (B) but for the percentage usage of isoform ENST00000449768 of OAS2. In all the panels NLS are highlighted by blue dots. The arrows on panel A-C highlight the location of the SNPs for which the boxplots are shown on the right.

In addition to overall changes in expression, we took advantage of the power of RNA-sequencing data to test if NLS in the OAS regions influenced the ratio of alternative isoforms used for each of the OAS genes (i.e., alternative splicing QTL: asQTL). We found that SNPs associated with the Neandertal haplotype are significant asQTL for *OAS1* and *OAS2* in both infected and non-infected macrophages (FDR << 1%; Figure 4B-C). The effect of the splice site variant rs10774671 at determining what isoform is primarily encoded by *OAS1* was particularly strong (*P* ≤ 2×10^−32^). This SNP is also a strong asQTL and protein QTL in lymphoblastoid cell lines [34,35]. The ancestral G allele at this SNP (AG at acceptor site) retains the splice site whereas the derived allele, A, (AA at acceptor site) disrupts the splice site leading to the usage of a distinct isoform (Figure S6). The Neandertal haplotype harbors the ancestral allele (encoding the p46 isoform), which is associated with high enzyme activity [36]. Interestingly, despite the fact that this ancestral allele is found at ~ 60-70% in most Sub-Saharan African populations, outside of Africa this allele is only found on individuals with the Neanderthal haplotype (with rare exceptions; ~2% of all haplotypes, Figure S2 suggesting that the Neandertal segment is effectively reintroducing an ancestral variant that was already present in other human groups. To validate that this SNP had the same impact at determining what *OAS1* isoforms are expressed in African-descent individuals, we analyzed additional RNA-sequencing data collected from 41 African-American individuals. As among Europeans, rs10774671 is a strong asQTL for *OAS1* in both non-infected and infected macrophages (P < 1×10^−15^, Figure S7), despite laying in a non-Neandertal haplotype.

Because OAS genes are primarily involved in the control of viral infections we decided to validate our functional findings on peripheral blood mononuclear cells (PBMCs) from 40 individuals stimulated/infected with viral-ligands (polyI:C and gardiquimod), and live viruses (Influenza, Herpes simplex virus (HSV) 1 and HSV2). The individuals were chosen based on their genotype for the NLS rs1557866, a SNP that is a strong proxy for the presence or absence of the Neandertal haplotype in the OAS region (9 were homozygous for the Neandertal haplotype, 15 were heterozygous, and 16 homozygous for the modern human sequence).

As expected, we found that all viral-associated immune triggers led to a marked increase in *OAS1-3* gene expression levels, as measured by real-time PCR (up to 27-fold, *P* ≤ 1.2×10^−8^, Figure 5A), concomitantly with the up-regulation of type-I and type-II interferon genes (Figure S8). Confirming the QTL results obtained in macrophages, we found that rs10774671 was a strong asQTL for *OAS1* in both non-infected and infected PBMCs (*P* ≤ 4.9×10^−5^, Figure 5B). Likewise, we found that the presence of the Neandertal haplotype was associated with reduced expression levels of *OAS3,* particularly in PBMCs infected with influenza (P = 6.1×10^−3^) and the synthetic ligand gardiquimod (P =2.0×10^−4^), which mimics a single strand RNA infection (Figure 5B). Interestingly, the Neandertal haplotype harbors additional regulatory variants that only impact expression levels in a cell-type and immune stimuli specific fashion. For example, we found that the Neandertal haplotype is associated with increased expression levels of *OAS2* in non-infected (*P* = 1.2×10^−3^), and gardiquimod-stimulated PBMCs (*P* = 4.9×10^−3^), but neither in macrophages nor in PBMCs treated with other viral agents. Collectively, our functional data provide evidence for a pervasive impact of the Neandertal haplotype on the regulation of *OAS* genes that varies depending on the cell type and the immune stimuli to which the cells are responding.

**Figure 5.**
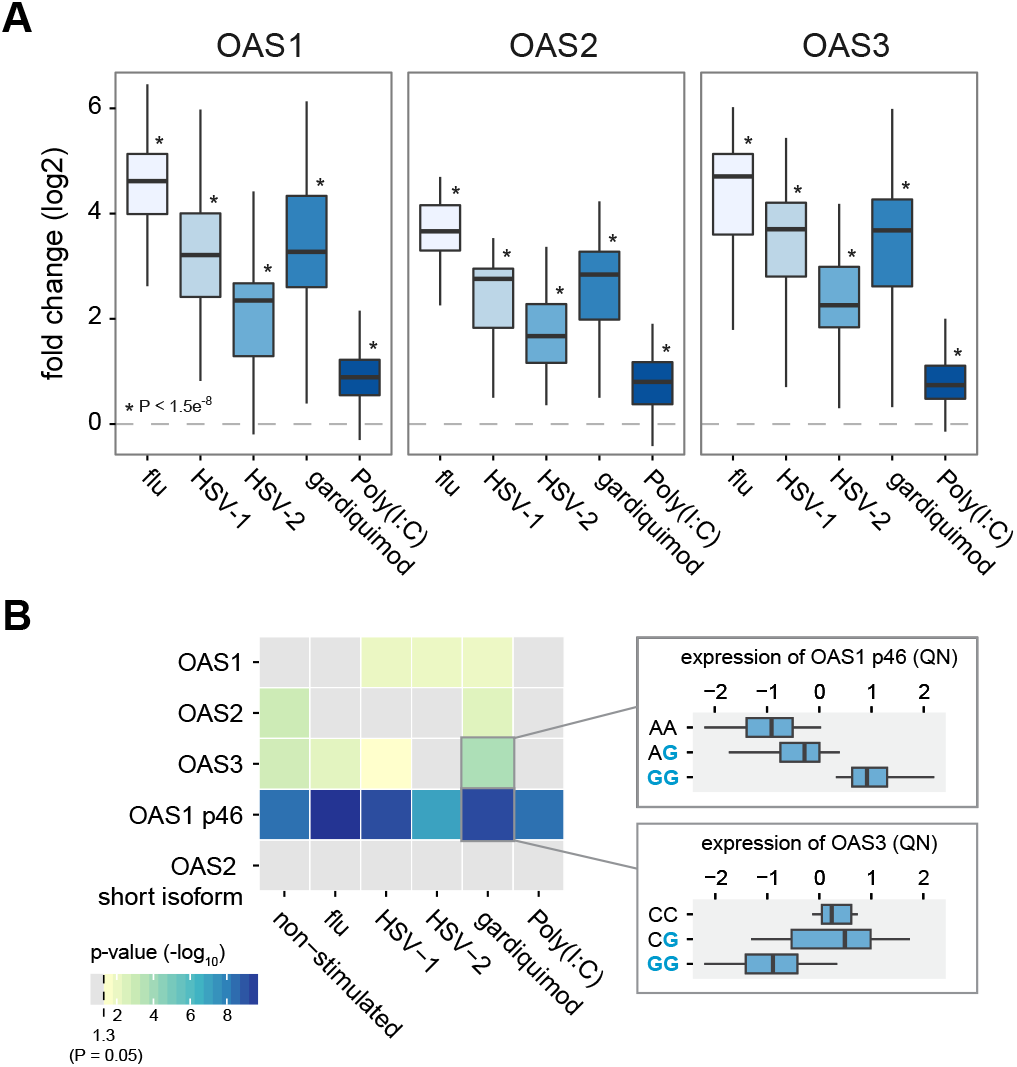
The Neandertal haplotype in the OAS regions has a different impact on the regulation of OAS genes depending on the viral agents PBMCs are exposed to. (A) Log 2 fold induction (y-axis) of *OAS1*, *OAS2* and *OAS3* in response to different viral agents or viral-associated immune stimuli, relative to non-infected PBMCs (B) -log10 P for the association between genotype status for the Neandertal haplotype and overall expression levels of OAS genes and the expression of specific isoforms of *OAS1* and *OAS2* (those identified in Figure 3 as associated with NLS). Boxplots show the association between genotype status (blue referring to Neanderthal alleles) and expression levels of the p46 isoform of *OAS1* and *OAS3* in Gardiquimod-stimulated PBMCs.

## Discussion

The Neandertal lineage was present in Eurasia for at least 400,000 years [37], providing ample time for Neandertals to adapt to local disease environments. The admixture process, which likely fostered the transmission of pathogens between Neandertals and humans migrating out of Africa, could have also led to the exchange of genes useful in responding to local pathogens. Here, we have demonstrated that a previously reported case of Neandertal introgression at the OAS locus [15] displays signatures of positive selection in the European population. Additionally, we have strengthened the case for adaptive introgression by providing direct functional evidence of a role for the Neandertal OAS haplotype in the regulatory responses in innate immune cells to infectious agents.

Our results show that the Neandertal haplotype at OAS is associated with several regulatory variants that reduce expression of *OAS3* in response to infection, as well as encode alternate isoforms of *OAS1* and *OAS2*. These dramatic functional implications of the Neandertal OAS haplotype support our case for adaptive introgression at OAS. Yet, because distinct functional polymorphisms segregate together in the same haplotype, inferring the exact variant(s) targeted by positive selection remains a daunting task. We speculate, however, that one of the strongest direct targets of selection is likely to have been the splice variant identified in *OAS1*.

The Neandertal haplotype carries the ancestral allele (G) of the *OAS1* splice variant (rs10774671), which is common both inside and outside of Africa. However, outside of Africa, the only haplotypes carrying this ancestral splice site are closely related to the Neandertal haplotype, with a few exceptions being rare recombined haplotypes (~2% of all haplotypes with the ancestral allele). This pattern reflects the possibility that Neandertal introgression, in effect, served as a means to resurrect the ancestral splice site from local extinction outside of Africa, probably following the out-of-Africa exodus. Interestingly, the same ancestral splice site is found in the Denosivan genome, and may similarly have impacted adaptive introgression into the ancestors of Melanesians. While we cannot at present confirm that the ancestral splice site was missing from the ancestral Eurasian population, the presence of this allele only on the Neandertal haplotype hints at the possibility that this splice site was lost to drift and subsequently re-introduced by Neandertals, providing beneficial genetic variation at the OAS locus (Figure S2). The Neandertal-introgressed allele encodes a protein variant (p46) that is associated with higher enzymatic activity [36]. The adaptive potential of this variant is supported by the observation that this variant (or other variants in strong LD with it) was shown to be associated with: *(i)* reduced infection and replication rates of West Nile virus ([38], but see [39]), *(ii)* improved resistance to hepatitis C virus (HCV) infection [40,41], and *(iii)* variable symptomology of Tick-Borne Encephalitis (TBE) Virus-Induced Disease (homozygous individuals for the Neandertal haplotype show the most severe symptoms of TBE). Strikingly, West Nile, hepatitis C and TBE are all members of the *Flaviviridae* family, suggesting that Flaviviruses might have been the main drivers of selection in *OAS1*.

The differential responses of homozygous carriers of the Neandertal OAS haplotype to the different viruses described above suggest that the Neandertal haplotype is not uniformly beneficial in humans. Thus, it is plausible that both spatial and temporal fluctuations in virus populations could have led to fluctuating selection pressure on Neandertal OAS haplotypes, consistent with findings that genes in the immune system have been disproportionately targeted by positive selection since the dawn of agriculture [42].

Alternatively, and not mutually exclusively, alleles at OAS, and particularly the *OAS1* splice variant, might be evolving under balancing selection. This hypothesis is supported by the observation that the *OAS1* splice variant (rs10774671) is found at high frequency worldwide (0.11-0.7), and the significantly higher Tajima’s D values observed around *OAS1*, as compared to genome-wide expectations (Figure 3D). Moreover, *OAS1* is among the most diverse genes in both humans and non-human primates. Indeed, a recent analysis of genome-wide sequence data from a total of 55 individuals from four non-human ape species, chimpanzee (*Pan troglodytes ellioti*), bonobo (*Pan paniscus*), gorilla (*Gorilla gorilla gorilla*), and orangutan (*Pongo abelii*), identified *OAS1* as in the top 1% of genes showing the largest levels of nucleotide diversity among ape species, consistent with a scenario of long-term balancing selection (*OAS2* and *OAS3* are ranked in the 60th and 36th percentile of the genome-wide distribution, respectively) or, as previous research has suggested, rapid evolution across the primate order [43]. Further supporting the idea of balancing selection on the introgressed haplotypes, our functional data suggest that the Neandertal haplotype contributes a range of gene expression responses in a cell-type and stimulus-specific manner.

Estimates of historical allele frequencies at the OAS locus support the notion that the Neandertal haplotype did not follow a classic selective sweep model with constant directional selection. Under such a model, the observed present-day frequency of the Neandertal haplotype at the OAS locus of roughly 38% would suggest a codominant fitness effect of *s* ~ 0.0014 − 0.0017, when assuming an initial frequency of 0.02 at introgression approximately 2,000-2,400 generations ago (Methods). However, the observed allele frequency shift of 0.26 over the past 200-340 generations (maximum shift in CEU from ancient samples; see Figure 2B) suggests that the selection coefficient associated with the Neandertal haplotype during this recent human evolution would have been *s* ~ 0.0044 −0.0075: 2.6−5.4 times larger than the above estimate. Therefore, both temporally varying selection and balancing selection could explain why the Neandertal haplotype at OAS did not show clear signatures of adaptive introgression in previous studies [15,18].

Our study illustrates the difficulty of identifying strong candidates for adaptive introgression. In a genome-wide statistical framework, our use of neutral coalescent simulations as a null distribution for adaptive introgression likely would not have provided significant results after multiple test correction across loci. This suggests that other instances of less obvious adaptive introgression, especially those that did not follow a classic selective sweep model, may remain to be identified. Moving forward, novel methods must be developed to identify such cases. Some progress is now being made on this front. For example, Racimo and colleagues [11] recently developed a genome-wide statistical framework which relies on distributions of uniquely shared derived alleles between humans and Neandertals (as does our study) to identify candidate regions for adaptive introgression. This study also highlighted the OAS locus as a candidate for adaptive introgression. Additionally, estimates of historical allele frequencies with increased spatial and temporal resolution provided by the sequencing of ancient human genomes are likely to play an important role in illuminating candidates for adaptive introgression which do not conform to classical selective sweep models.

## Conclusions

In conclusion, our study demonstrates that the frequency and haplotype distribution of Neandertal-like sites can be used in a neutral simulation framework that accounts for local genomic context to investigate the history of selection at a candidate locus for which genome-wide tests of selection provide ambiguous results. When combined with functional data, our results provide the strongest evidence to date in support of adaptive introgression in the OAS region. More generally, our study raises the possibility that adaptive introgression might not necessarily occur to select newly introduced variants but rather as a means to resurrect adaptive variation in modern human populations that had been lost due to demographic events.

## Materials and Methods

### 1. Genome alignments and identification of Neandertal-like sites

Human/chimpanzee ancestral states were computed by parsimony using alignments from the UCSC Genome Browser for the human reference (hg19) and three outgroups chimpanzee (panTro2), orangutan (ponAbe2), and rhesus macaque (rheMac2) [44]. Ancestral state was assumed to be the chimpanzee allele (if available) if its state was confirmed by matching either orangutan or macaque. All sites with no inferred ancestral state were removed from our analysis.

We filtered the Altai Neandertal genome [4] and Denisovan genome [3] using the map35_50 set of minimum filters provided at (https://bioinf.eva.mpg.de/altai_minimal_filters/). For the frequency analysis, we combined these filtered datasets with ten samples from outside of Africa (5-Eur European: CEU, FIN, GBR, IBS, TSI; 5-East Asian: CDX, CHB, CHS, JPT, KHB; see Table S7 for population codes) and one sub-Saharan African sample YRI (Yoruba in Ibadan, Nigeria) from the 1000 Genomes Project Phase 3 [20], which we downloaded from (https://mathgen.stats.ox.ac.uk/impute/1000GP%20Phase%203%20haplotypes%206%20October%202014.html).

We first extracted all biallelic variants in our human, Neandertal, and chimpanzee alignments. We considered as Neandertal-like sites (NLS) only those variants where the African sample (YRI) had a derived allele frequency of zero and both the non-African sample and Neandertal carried the derived allele. We restricted the follow-up haplotype based analysis to only the CEU sample. In that analysis we required the derived allele to be present in CEU in at least two copies in order to calculate our haplotype-based test statistic (*H*_*D/A*_).

### 2. Haplotype clustering in the 1000 Genomes Project data

To assess the relationships and frequencies of Neandertal-like haplotypes in the OAS region, we identified haplotype clusters based on sequence similarity. First, we filtered all 5,008 haplotypes in the 1000 Genomes Project phase 3 dataset to include only SNPs within the bounds of the three OAS genes (hg19 chr12: 113344739-113449528) and at which the Altai Neandertal [4]and Denisovan [3] genomes carry homozygous genotypes. Next, we clustered the human haplotypes such that all haplotypes in a cluster had no more than 60 pairwise differences, which resulted in 10 clusters. We inferred a consensus haplotype for each cluster based on the majority allele at each position in the cluster. Finally, we produced a neighbor-joining tree of the human cluster consensus sequences and the Altai, Denisovan, and ancestral sequences using the R package “ape.” We estimated confidence values for nodes with one thousand bootstraps.

We then calculated haplotype frequencies for each of the 1000 Genomes Project population samples based on the assignment of each haplotype to the clusters described above.

### 3. Demographic model and neutral coalescent simulations

We performed coalescent simulations of the demographic history of the European, East Asian, African, and Neandertal populations applied by Vernot and Akey [25] based on previously inferred demographic models [23,24]. In our first set of simulations, which were performed with ms [45], we extracted only allele frequencies at NLS. We gathered a total of 1 million simulation results. In each simulation run, a result was collected if a NLS was present in either the European or the East Asian population sample (or both). Therefore, null distributions specific to Europeans or East Asians included less than 1 million results (but typically > 800,000). Haplotype simulations were performed with macs [46] in order to explicitly simulate the genetic map (downloaded with the 1000 Genomes samples at the link above) of the 600 kilobase region centered on OAS (chr12:113100000-113700000). The general shape of the demographic model is illustrated in Figure S1. Our baseline simulations were performed with the parameters specified in Table S1, assuming 25 years per generation and a mutation rate of 2.5 × 10^−8^ per bp per generation. Table S1 also provides all parameters used in one- and two- introgression pulse simulations. Additionally, we examined the one-pulse model with a slower mutation rate (1.25 × 10^−8^) and a one-pulse model with a uniform recombination rate (1.3 × 10^−8^; a value conservatively lower than the average for the OAS region, 1.7 × 10^−8^). Sample ms and macs commands are given at the end of this section.

For haplotype-based simulations, data were thinned in a manner similar to Sankararaman *et al.* 2014 [6] to account for imperfect SNP ascertainment in the 1000 Genomes dataset, such that SNPs with minor allele count of 1, 2, 3, 4, 5, 6, 7, 8, 9, and > = 10 were accepted with probabilities 0.25, 0.5, 0.75, 0.8, 0.9, 0.95, 0.96, 0.97, 0.98, and 0.99, respectively. Additionally, we only kept SNPs that were polymorphic in the simulated CEU sample. Finally, we performed an additional thinning of SNPs with uniform probability of 0.05 of removal to account for slightly elevated SNP density in the simulated data. The resulting simulated datasets had an average SNP density of 3.8 SNPs per kb compared to 2.5 in the real data. This is a slightly larger than ideal difference in SNP density, but we note that neither derived allele frequency, nor our primary haplotype-based test statistic (described below) should be particularly sensitive to SNP density. In fact, Figure S9 illustrates that our statistic is conservative with respect to SNP density. Additionally, our results hold in a simulation with half the mutation rate of our baseline simulation. This simulation resulted in an average SNP density of 1.9 SNPs per kb. So our observation of long haplotypes surrounding NLS is robust to SNP density across our simulations.

#### Sample ms command

ms 752 1 -s 1 -I 4 250 250 250 2 0 -n 1 58.0027359781 -n 2 70.0410396717 -n 3 187.549931601 -n 4 0.205198358413 -eg 0 1 482.67144247 -eg 0 2 505.592963281 -eg 0 3 720.224280773 -em 0 1 2 0.358619661426 -em 0 2 1 0.358619661426 -em 0 1 3 0.111889334365 -em 0 3 1 0.111889334365 -em 0 2 3 0.446122858814 -em 0 3 2 0.446122858814 -en 0.00699726402189 1 1.98002735978 -eg 0.00699726402189 2 0.0 -eg 0.00699726402189 3 17.5076517287 -en 0.03146374829 2 2.03666547538 -en 0.03146374829 3 0.700185007205 -ej 0.0641347992705 3 2 -em 0.0641347992705 1 2 0.00015 -em 0.0641347992705 2 1 0.00015 -em 0.0839811779742 2 4 0.00075 -em 0.0846651725023 2 4 0 -ej 0.0957592339261 2 1 -en 0.202462380301 1 1.0 -ej 0.957592339261 4 1

#### Sample macs command

macs 416 600000 -t 0.000731 -R oas_recrates.txt -I 4 216 198 0 2 0 -n 1 58.0027359781 -n 2 70.0410396717 -n 3 187.549931601 -n 4 0.205198358413 -eg 0 1 482.67144247 -eg 1e-12 2 460.409556336 -eg 2e-12 3 720.224280773 -em 3e-12 1 2 0.4441436762 -em 4e-12 2 1 0.4441436762 -em 5e-12 1 3 0.138572826974 -em 6e-12 3 1 0.138572826974 -em 7e-12 2 3 0.552514733193 -em 8e-12 3 2 0.552514733193 -en 0.00699726402189 1 1.98002735978 -eg 0.00699727402189 2 0.0 -eg 0.00699728402189 3 18.896348561 -en 0.03146374829 2 2.79399962717 -en 0.03146375829 3 0.670250141711 -ej 0.051785044418 3 2 -em 0.051785054418 1 2 0.00015 -em 0.051785064418 2 1 0.00015 -em 0.0555806720117 2 4 0.00075 -em 0.0562646665398 2 4 0 -ej 0.0957592339261 2 1 -en 0.202462380301 1 1.0 -ej 0.957592339261 4 1 -h 1e3

### 4. Frequency and haplotype-based tests of neutrality using simulations

We examined the consistency of genetic variation with our neutral model using several approaches. First, we examined the likelihood of observing (Neandertal) allele frequencies as high as the OAS locus. Under neutrality, allele frequency is not dependent upon recombination rate, therefore, we can estimate the likelihood of our observed NLS frequency in the OAS region. To create a distribution of expected NLS frequency in each introgression model, we first extracted derived allele counts (DAC) at simulated NLS for samples of 250 individuals each for Europeans and East Asians in the simulated model. Next, for each non-African population sample from the 1000 Genomes Project, we binomially sampled the DACs from the appropriate simulated population (European for CEU, GBR, IBS, TSI; East Asian for CDX, CHB, CHS, JPT, KHV) using the true population sample size to create a distribution of NLS frequencies that is specific to each population sample (Figure 2A, S3). Minimum p-values across the OAS locus for each population are listed in Table S2.

Additionally, we wanted to examine if the haplotypes carrying NLS at the OAS locus are longer than expected under neutrality when conditioning on the observed frequencies and the underlying genetic map, which would provide an additional signature of selection on introgressed haplotypes beyond empirical haplotype signatures. For this purpose, we modified a simple haplotype statistic *H* [47], which measures the average length of pairwise homozygosity tracts in base pairs – a quantity that is very straightforward to interpret. As selective sweeps are expected to create long haplotypes around the selected site, the *H* statistic should be higher in samples containing positively selected haplotypes compared to samples containing neutrally evolving haplotypes, when frequency and recombination are properly controlled, similar to other statistics based on haplotype lengths, such as *EHH, iHS,* and *nSL* [28,30,48]. However, in contrast to these other statistics, *H* does not require specification of analysis parameters such as minimum haplotype homozygosity levels below which haplotypes are no longer extended.

Under adaptive introgression, we specifically expect the introgressed (derived allele carrying) haplotypes to be longer than the ancestral haplotypes, due to the fact that selected Neandertal haplotypes will on average spend less time at low frequency, where they have a greater opportunity for recombination with non-introgressed haplotypes, than neutral Neandertal haplotypes. We therefore defined our test statistic, *H*_*D/A*_, as:

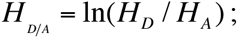

where *H*_*D*_ and *H*_*A*_ are *H* calculated across haplotypes carrying derived NLS allele versus the ancestral allele (see Figure S10).

For each model we generated 1,000 simulated OAS regions containing NLS that reach a frequency greater than the 99^th^ percentile for CEU in our frequency simulations. We calculated *H*_*D/A*_ for every simulated NLS and for every NLS in our true sample. We then compared our simulation results to the observed data in two ways. First, to visually examine the distribution of true *H*_*D/A*_ values to neutral simulations, we took the mean *H*_*D/A*_ score across all NLS in each of 1,000 simulations and calculated the 95^th^ and 99^th^ percentiles of those 1,000 simulations (Figure 3B). Second, to examine the probability under neutrality of the observed *H*_*D/A*_ values at the peak of *H*_*D/A*_ between chr12:113380000-113420000, we compared the mean *H*_*D/A*_ score for SNPs in this window to the mean of SNPs in this window for each simulation that produced a NLS in that window. These results for all tested models are listed in Table S4.

### 5. Analysis of ancient Eurasian data

We utilized supplementary data table 3 from Mathieson *et al.* [26]. This table includes maximum likelihood allele frequency estimates for three ancient population samples (HG- Hunter-gatherer, EF- Early farmer, SA- Steppe ancestry) and four present day European samples from the 1,000 Genomes Project (see “Genome-wide scan for selection” section of methods in[26]). We intersect this table with allele frequencies for 1,000 Genomes Yorubans (YRI) and the Altai Neandertal genotypes and only analyze sites for which we have data for all samples (1,004,612 SNPs).

To calculate the expected allele frequency in modern samples under drift, we used the estimated proportions (*m*) of (HG, EF, SA) in each of the four present-day samples estimated by Mathieson *et al.* [26]: CEU = (0.196, 0.257, 0.547), GBR = (0.362, 0.229, 0.409), IBS = (0, 0.686, 0.314) and TSI = (0, 0.645, 0.355). We calculated the expected frequency E[*p*] of site as:

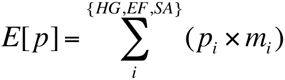

Next, we calculated the absolute difference between observed and expected allele frequency in all four present-day European samples at all available sites. To test the null hypothesis that OAS NLS have not changed in frequency more than expected under neutrality at 11 SNPs in the central OAS region (chr12:113200000-113600000), we first calculated the fraction of all autosomal SNPs in the dataset at similar present-day frequency (within 1 percent in the folded frequency spectrum) of each OAS NLS. Finally, we calculated the fraction of autosomal SNPs with an absolute observed minus expected frequency difference greater than or equal to the OAS NLS. These results are given in Table S3 and illustrated in Figure 2B.

This test does not explicitly incorporate variance in estimated ancient allele frequency. However, any bias in ancient allele frequency estimation should be distributed randomly across the genome. Therefore, our comparison to a genome-wide distribution of SNPs at similar present-day frequency should incorporate most of this error. Nonetheless, the selection test performed by Mathieson *et al.* [26] does incorporate such error, so we can also look to the Ps from that test to ensure consistency with our results.

### 6. Estimation of selection coefficients

To estimate the selection coefficient *s* under constant positive selection for a given starting frequency (*x*_*0*_), final frequency (*x*_*1*_), and number of generations between these estimates (*Δt*), we assumed a model of standard logistic growth of a codominant allele:

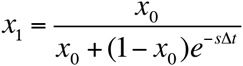

This equation can be easily solved to obtain *s*, given *x*_*0*_, *x*_*1*_, and *Δt*.

### 7. Probabilities that Neandertal haplotypes are shared due to incomplete lineage sorting

We estimated the probability that Neandertal haplotypes observed in the OAS region could be a product of ILS using the method outlined by Huerta-Sanchez *et al.* [49]. This method estimates the probability that a haplotype of length (H) is shared due to incomplete lineage sorting as:

*P(H) = 1 - gammaCDF(H, shape = 2, rate = 1/L);*

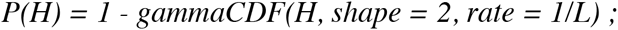

where L is the expected shared length, L = 1/(r × t) where r is recombination rate and t is time in generations.

We first estimated the maximum haplotype length observed in each of the 1000 Genomes Project population samples by identifying tracts of identity-by-state (IBS) that include NLS between the Altai Neandertal haplotype (considering only homozygous positions) and individual human haplotypes. We identified all tracts of IBS bounded by NLS (using YRI as the African reference) and took the maximum of these per-population sample IBS distributions.

We identified the maximum tract length (198,441 bp) in European (and some admixed Native American) samples. Maximum tract length for East Asian samples (54,385 bp) is notably shorter. As expected, no IBS tracts were identified in African population samples.

Assuming a conservatively low recombination rate for the OAS region (1.3 × 10^−8^) and a conservatively short divergence between Neandertals and humans of 300 thousand years (and 25 years per generation), the probability of a length of at least 54,385 is P(54385) = 0.002. The probability of the longest tracts is much smaller, P(198441) = 1.15 × 10^−12^. Therefore, it is unlikely that Neandertal haplotypes are shared with non-Africans due to shared ancestral variation.

We repeated this analysis using the Denisovan genome and found no IBS tracts defined by Denisovan-like sites, suggesting that introgressed Denisovan haplotypes described by Mendez *et al.* [17] are not present in the 1000 Genomes Project Samples, consistent with our findings in Figure 1A.

### 8. Neutrality tests in phase 3 of the 1000 Genomes data

We calculated iHS, DIND, Fst and Tajima’s D on different populations from phase 3 of the 1000 Genomes project. The phased data were downloaded from Impute reference data panel (https://mathgen.stats.ox.ac.uk/impute/1000GP_Phase3.html) and were filtered for biallelic SNPs. Arlequin version 3.5.1.3 [50] was then used to calculate Fst estimates derived from ANOVA among all pairs of European and Asian populations. DIND was calculated as previously described [29] and Tajima’s D was calculated for all SNPs using a window of 25kb either side of each SNP (i.e., in window of 50kb in total). iHS values were recovered from a genomic scan performed using hapbin [51] on the same phase 3 release of the 1000 Genomes dataset.

### 9. Sample collection

Buffy coats from 96 healthy European American donors and 41 African Americans were obtained from Indiana Blood Center (Indianapolis, IN, USA). Only individuals self-reported as currently healthy and not under medication were included in the study. The project was approved by the ethics committee at the CHU Sainte-Justine (protocol #4022). The individuals recruited in this study were males aged 18 to 55 years old. For 80% of our samples we have information on their exact age (for the remaining 20% of donors we only know that they were adults less than 55 years old), and we found no association between the presence of the Neandertal haplotype (as measure by rs1557866) and age (*P* > 0.5). Thus, variation in age is not likely impact any of our conclusions. In addition to self-identifed ancestry labels, we used the genome-wide genotype data (see section 10) to estimate genome-wide levels of European and African ancestry in each sample using the program ADMIXTURE [52]. Consistent with previous reports, we found that self-identified EA showed limited levels of African admixture (mean = 0.4%, range 0-13%). The African Americans used in our study were selected on having at least 75% of African ancestry.

### 10. DNA Extraction and genotyping

DNA from each of the blood donors was extracted using the Gentra Pure Gene blood kit (Qiagen). Genotyping of each individual was then performed by Illumina’s HumanOmni5Exome bead Chip array and complemented with imputed data from the 1000 Genomes data using Impute2 [53]. Here, we only focused on genetic diversity surrounding the OAS region – chr12:113229549-113574044 (~ 344Kb) spanning from the beginning of *RPH3A* to the end of *RASAL1* – for a total of 673 SNPs with a MAF above 10%. Genotypes can be found in Table S8.

### 11. Isolation of monocytes and differentiation of macrophages

Blood mononuclear cells were isolated by Ficoll-Paque centrifugation. Monocytes were purified from peripheral blood mononuclear cells (PBMCs) by positive selection with magnetic CD14 MicroBeads (Miltenyi Biotech) using the autoMACS Pro Separator. All samples had purity levels above 90%, as measured by flow cytometry using an antibody against CD14 (BD Biosciences). Monocytes were then cultured for 7 days in RPMI-1640 (Fisher) supplemented with 10% heat-inactivated FBS (FBS premium, US origin, Wisent), L-glutamine (Fisher) and M-CSF (20ng/mL; R&D systems). Cell cultures were fed every 2 days with complete medium supplemented with the cytokines previously mentioned. Before infection, we systematically verified that the differentiated macrophages presented the expected phenotype for non-activated macrophages (CD1a+, CD14+, CD83–, and HLA-DRlow (BD Biosciences)).

### 12. Bacterial preparation and infection of macrophages

The day prior to infection, aliquots of *Salmonella typhimurium* (Keller strain) were thawed and bacteria were grown overnight in Tryptic Soy Broth (TSB) medium. Bacterial culture was diluted to mid-log phase prior to infection and supernatant density was checked at OD600. Monocyte-derived macrophages were then infected with *Salmonella typhimurium* at a multiplicity of infection (MOI) of 10:1. A control group of non-infected macrophages was treated the same way but using medium without bacteria. After 2 hours in contact with the bacteria, macrophages were washed and cultured for another hour in the presence of 50 mg/ml of gentamycin in order to kill all extracellular bacteria present in the medium. The cells were then washed a second time and cultured in complete medium with 3 mg/ml gentamycin for an additional 2 hours, the time point to which we refer in the main text.

### 13. Infection/stimulation of PBMC

PBMCs from a subset of 40 individuals used to derive macrophages were cultured in RPMI-1640 (Fisher) supplemented with 10% heat-inactivated FBS (FBS premium, US origin, Wisent) and 1% L-glutamine (Fisher). The 30 individuals were chosen based on their genotype for rs1557866, a SNP which derived allele is of Neandertal origin and that we used as a proxy to identify individuals harbouring the Neandertal haplotype in the OAS region. From the 40 individuals, and based on this SNP, 9 individuals were homozygous for the Neandertal haplotype, 15 were heterozygous, and 16 homozygous for the modern human sequence.

For each of the tested individuals, PBMCs (1 million per condition) were stimulated/infected with one of the following viral-associated immune challenges: polyI:C (10 μg/ml, TLR3 agonist), gardiquimod (0.5μg/ml, TLR7 and TLR8 agonist), Influenza PR8 WT (multiplicity of infection (MOI) of 0.05:1), Herpes simplex virus (HSV) 1 (1.55×10^2^ CPE), and HSV2 (19.5×10^4^ CPE). PBMCs were stimulated/infected for 4 hours with TLR ligands and Influenza, and 6h with HSV1 and HSV2. A control group of non-infected PBMC was treated the same way but with only medium.

### 14. RNA extraction, RNA-seq library preparation, and sequencing

Total RNA was extracted from the non-infected and infected/stimulated cells using the miRNeasy kit (Qiagen). RNA quantity was evaluated spectrophotometrically, and the quality was assessed with the Agilent 2100 Bioanalyzer (Agilent Technologies). Only samples with no evidence of RNA degradation (RNA integrity number > 8) were kept for further experiments. RNA-sequencing libraries were prepared using the Illumina TruSeq protocol. Once prepared, indexed cDNA libraries were pooled (6 libraries per pool) in equimolar amounts and sequenced with single-end 100bp reads on an Illumina HiSeq2500. Results based on the entire dataset are described elsewhere (Nédélec *et al.*, under revision). Here, we only studied transcript-level and gene-level expression estimates for *OAS1*, *OAS2* and *OAS3*. The transcript-level and gene-level expression data for these genes can be found in Table S9.

### 15. Quantifying gene expression values from RNA-seq data

Adaptor sequences and low quality score bases (Phred score < 20) were first trimmed using Trim Galore (version 0.2.7). The resulting reads were then mapped to the human genome reference sequence (Ensembl GRCh37 release 65) using TopHat (version 2.0.6) and using a hg19 transcript annotation database downloaded from UCSC. Gene-level expression estimates were calculated using featureCounts (version 1.4.6-p3) and transcript-level expression values were obtained using RSEM under default parameters.

### 16. Quantitative real time PCR

For the PBMC samples we measured the expression levels of OAS and interferon genes using real time PCR. 100ng of high-quality RNA was reverse-transcribed into cDNA using the qScript cDNA SuperMix (Quanta Biosciences). Quantitative real time PCR was performed using 96.96 Dynamic Array™ IFCs and the BioMark™ HD System from Fluidigm. For the TaqMan gene assays, we used the following TaqMan Gene Expression Assay (Applied BioSystems) to quantify the expression levels of interferon genes: *IFNA1* (Hs03044218), *IFNA6* (Hs00819627), and *IFNG* (Hs00989291). To quantify the overall expression levels of OAS genes, we used probes that capture all common isoforms of *OAS1* (Hs00973635), *OAS2* (Hs00942643), and OAS3 (Hs00196324). Custom-made probes were designed to specifically target the short-isoform of *OAS2* (Forward Primer Sequence CTGCAGGAACCCGAACAGTT; Reverse Primer Sequence ACTCATGGCCTAGAGGTTGCA; Reporter Sequence AGAGAAAAGCCAAAGAA). As housekeeping genes we used: *GAPDH* (Hs02758991), *GUSB* (Hs99999908), *HPRT1* (Hs99999909), and *POLR2A* (Hs00172187). The results reported in the manuscript used *POLR2A* as a reference but all conclusions remain unchanged when using any of the other housekeeping genes.

We start by doing a preamplification of the cDNA using the PreAmp Master Mix (Fluidigm). Preamplified cDNA was then diluted 2X on a solution of 10 mM Tris–HCl (pH 8.0) and 0.1 mM EDTA. In order to prepare samples for loading into the integrated fluid circuit (IFC), a mix was prepared consisting of 360 μL TaqMan Fast Advanced Master Mix (Applied BioSystems) and 36 μL 20× GE Sample Loading Reagent (Fluidigm). 2.75 μL of this mix was dispensed to each well of a 96-well assay plate and mixed with 2.25 μL of preamplified cDNA. Following priming of the IFC in the IFC Controller HX, 5 μL of the mixture of cDNA and loading reagent were dispensed in each of the sample inlet of the 96.96 IFC. For the TaqMan gene assays, 5 μL of mixes consisting of 2.5 μL 20× TaqMan Gene Expression Assay (Applied BioSystems) and 2.5 μL 2X Assay Loading Reagent (Fluidigm) were dispensed to each detector inlet of the 96.96 IFC. After loading the assays and samples into the IFC in the IFC Controller HX, the IFC was transferred to the BioMark HD and PCR was performed using the thermal protocol GE 96 × 96 Fast v1.pcl. This protocol consists of a Thermal Mix of 70 °C, 30 min; 25 °C, 10 min, Hot Start at 95 °C, 1 min, PCR Cycle of 35 cycles of (96 °C, 5 s; 60 °C, 20 s). Data was analysed using Fluidigm Real-Time PCR Analysis software using the Linear (Derivative) Baseline Correction Method and the Auto (Detectors) Ct Threshold Method.

To quantify the expression levels of the *OAS1* isoform associated with the derived allele at the splicing variant rs10774671 we used SybrGreen and the following forward (GCTGAGGCCTGGCTGAATTA), and reverse (CCACTTGTTAGCTGATGTCCTTGA) primers. PCR was performed using the thermal protocol 50 °C, 2 min; 95 °C, 10 min, PCR Cycle of 40 cycles of (95 °C, 15 s; 60 °C, 1 min). A melting curve was also performed to check for non-specific amplification.

### 17. Genotype–Phenotype Association Analysis

eQTL, asQTL were performed against *OAS1*, *OAS2* and *OAS3*. We examined associations between SNP genotypes and the phenotype of interest using a linear regression model, in which phenotype was regressed against genotype. In particular, expression levels were considered as the phenotype when searching for eQTL and the percentage usage of each isoform in each gene when mapping asQTL. To avoid low power caused by rare variants, only SNPs in the OAS region with a minor allele frequency of 10% across all individuals were tested (i.e., 673 SNPs within the region chr12:113229549-113574044). In all cases, we assumed that alleles affected the phenotype in an additive manner. For the eQTL and asQTL analyses on macrophages we mapped Salmonella-infected, and non-infected samples separately. We controlled for false discovery rates (FDR) using an approach analogous to that of Storey and Tibshirani[54], which makes no explicit distributional assumption for the null model but instead derives it empirically null from permutation tests, where expression levels were permuted 1000 times across individuals. For the non-infected and infected/stimulated PBMCs we only tested expression levels against the SNPs identified as eQTL or asQTL in the macrophage data (specifically, the SNPs for which boxplots are shown in Figure 4).

## Author Contributions

Conception and design: AJS, PWM, LBB; Acquisition of data: AJS, AD, YN, VY, LBB; Analysis and interpretation of data: AJS, AD, YN, PWM, LBB; Contributed unpublished, essential data, or reagents: CA, JET; Drafting or revising the article: AJS, PWM, LBB.

## Acknowledgements

We thank all members from the Barreiro and Messer labs for helpful discussions and comments on the manuscript. We thank Dr. Silvia Vidal for the gift of the Influenza PR8 WT used in this study, Dr. Hugo Soudeyns and Doris Ransy for advice with the viral infections, and Marc Montero and George (PJ) Perry for sharing their analysis on nucleotide diversity levels of OAS genes in non-human primate species.

## Availability of data and materials

The datasets supporting the conclusions of this article are included within the article and in supplementary Tables 8 and 9. The raw data were deposited under GEO accession numbers GSE73765 and GSE81046.

## Figures

**Figure S1.**
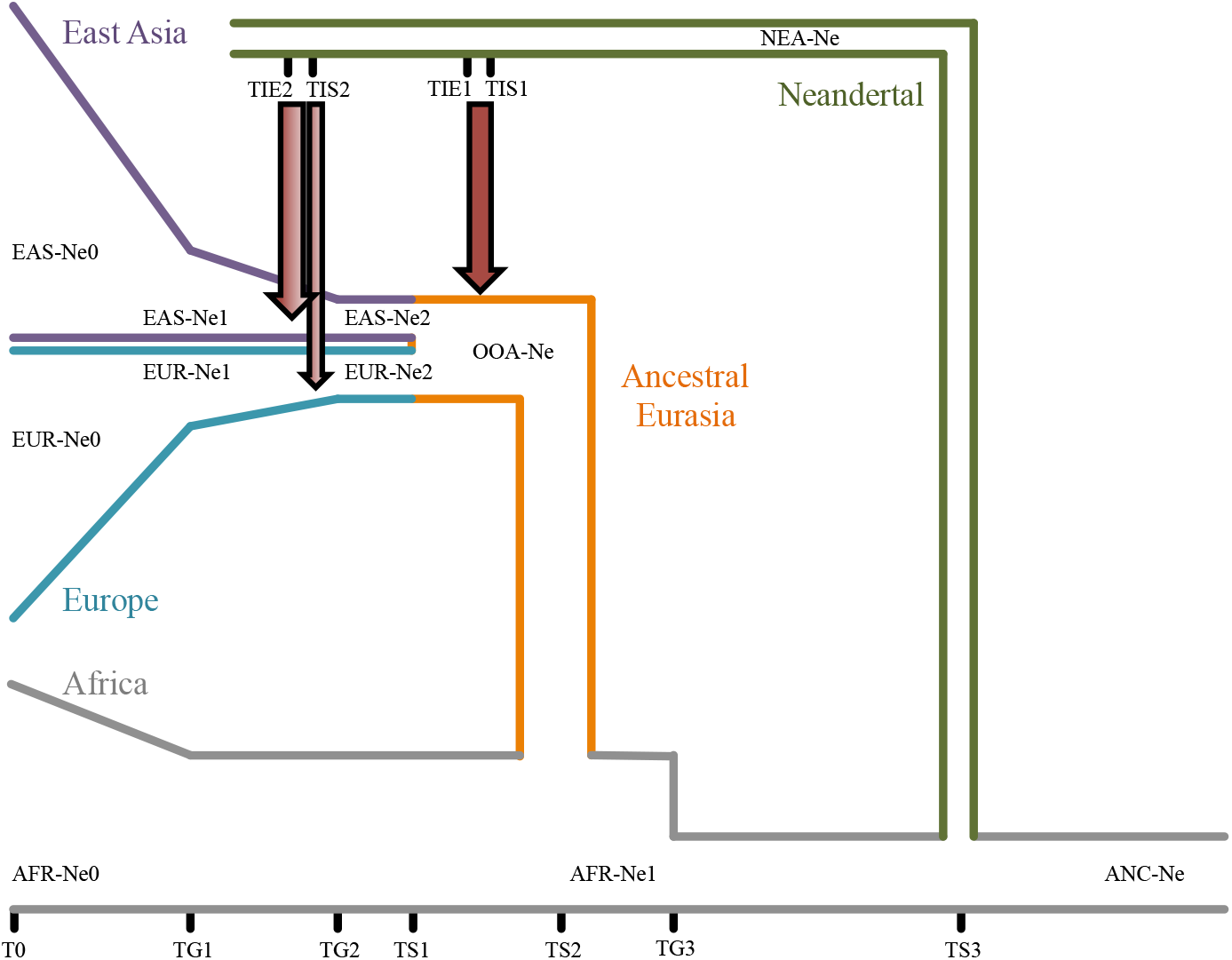
Schematic representation of demographic model used in simulations. Red arrows indicate points of introgression. The full range of demographic parameters used for the simulation can be found in Table S1.

**Figure S2.**
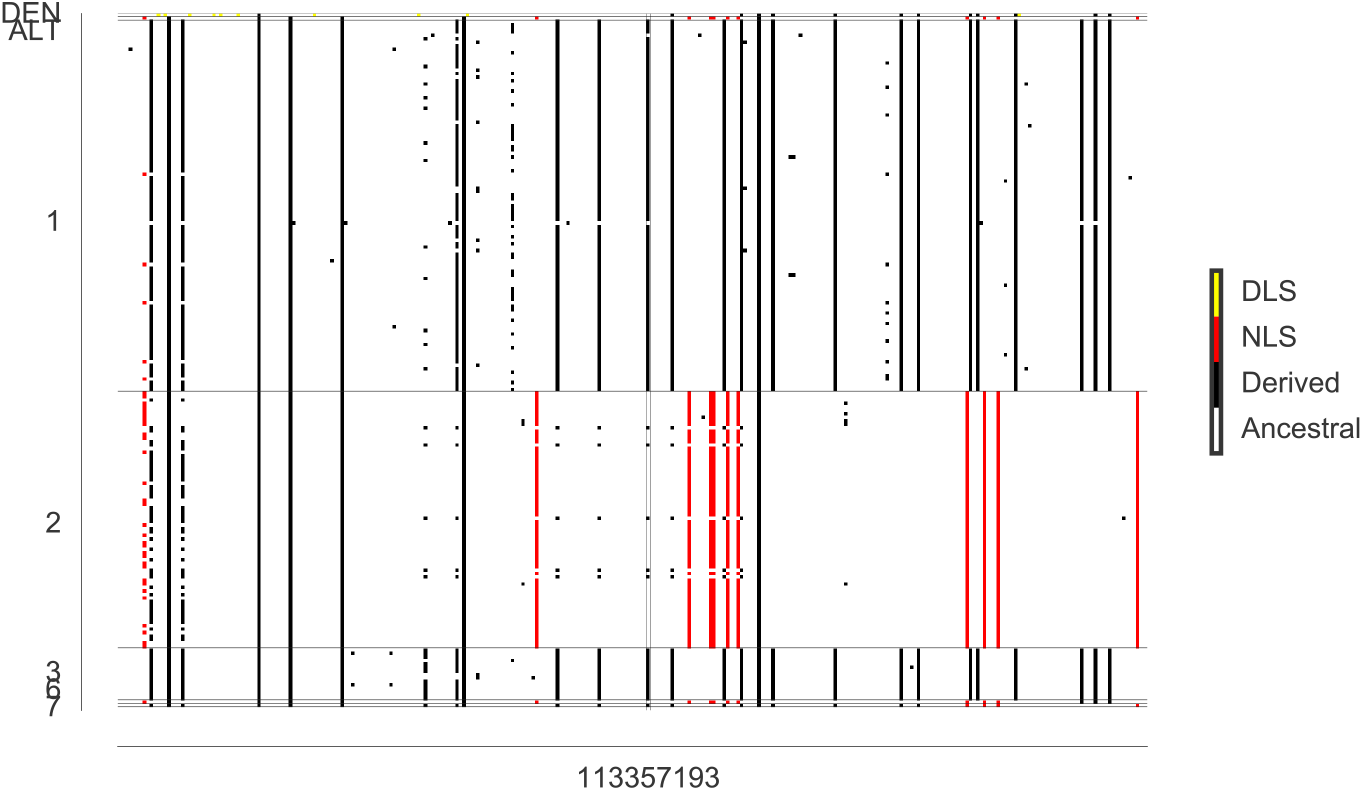
Haplogram of region surrounding *OAS1* splice site (rs10774671). 198 phased haplotypes from the CEU population sample are illustrated along with the Altai and Denisovan haplotypes. Ancestral alleles are colored white, derived alleles are (1) yellow if derived in Denisovan and absent from Africans (YRI) (DLS), (2) red if derived in Neandertals and absent from Africans (YRI) (NLS), (3) black otherwise. With the exception of two rare haplotypes in cluster 1, all occurrences of the ancestral variant at rs10774671 are surrounded by derived Neandertal alleles, indicating that these were introduced in a Neandertal haplotype. Only sites where the Neandertal and Denisovan genomes are homozygous were used in this haplogram. Cluster labels (y-axis) correspond to clusters in Figure 1.

**Figure S3.**
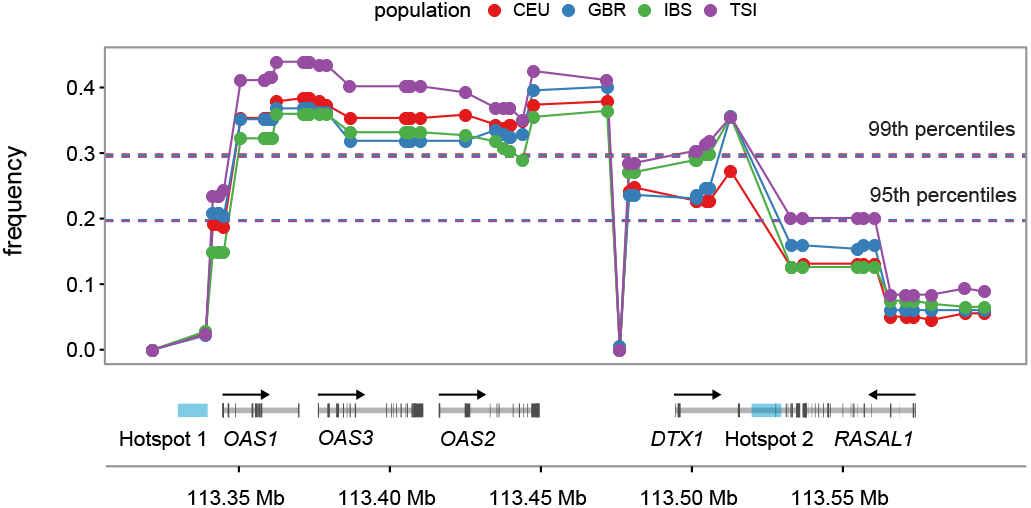
OAS-introgressed haplotypes are found at higher frequencies in European populations than expected under neutrality when using a two-pulse introgression model. (A) Comparison of frequency (y-axis) of NLS in the OAS locus in the CEU, GBR, IBS, and TSI European population samples with respect to neutral expectations (dashed lines) based on coalescent simulations considering two-pulse introgression in both Europeans and Asians, or a second pulse of introgression only in Europeans. Dashed lines overlap almost completely, reflecting little variation between a two-pulse introgression in both European and Asians, or a second pulse only in Europeans.

**Figure S4.**
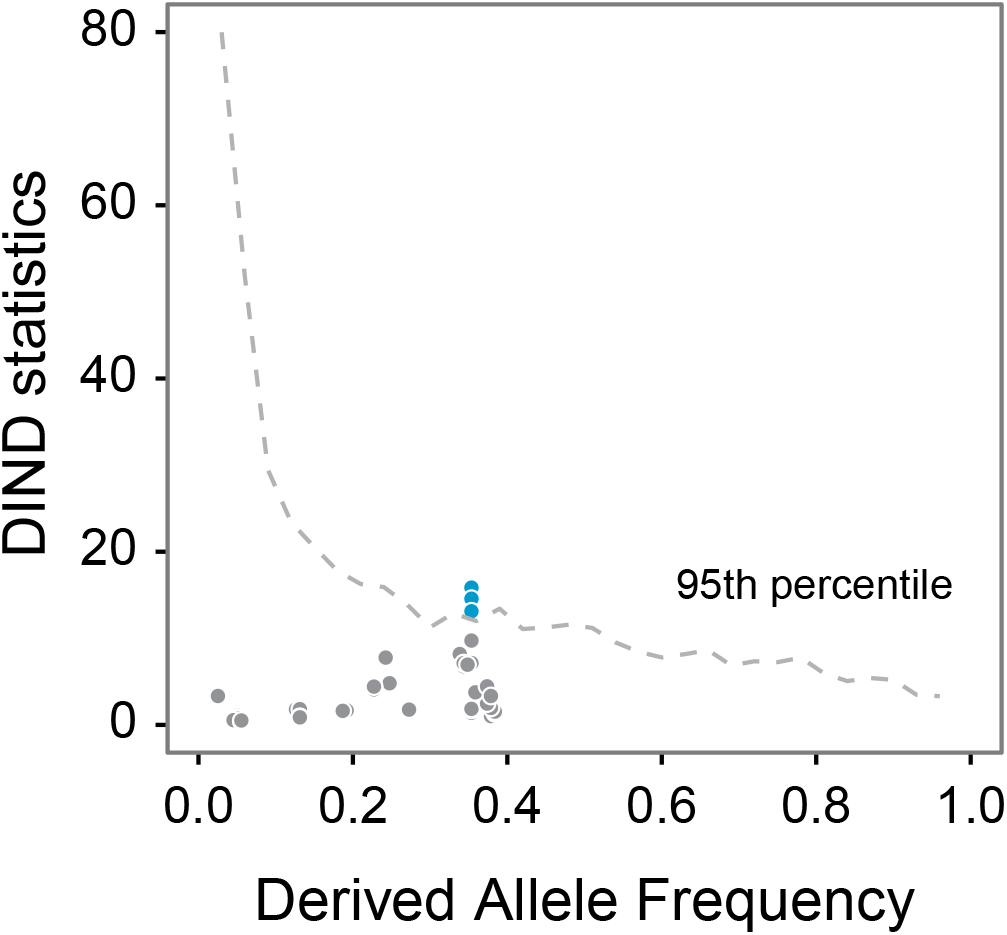
The DIND statistic indicates reduced diversity surrounding NLS. DIND (y-axis) is plotted for SNPs in OAS locus by derived (Neandertal) allele frequency. Dashed line indicates genome-wide 95^th^ percentile by frequency.

**Figure S5.**
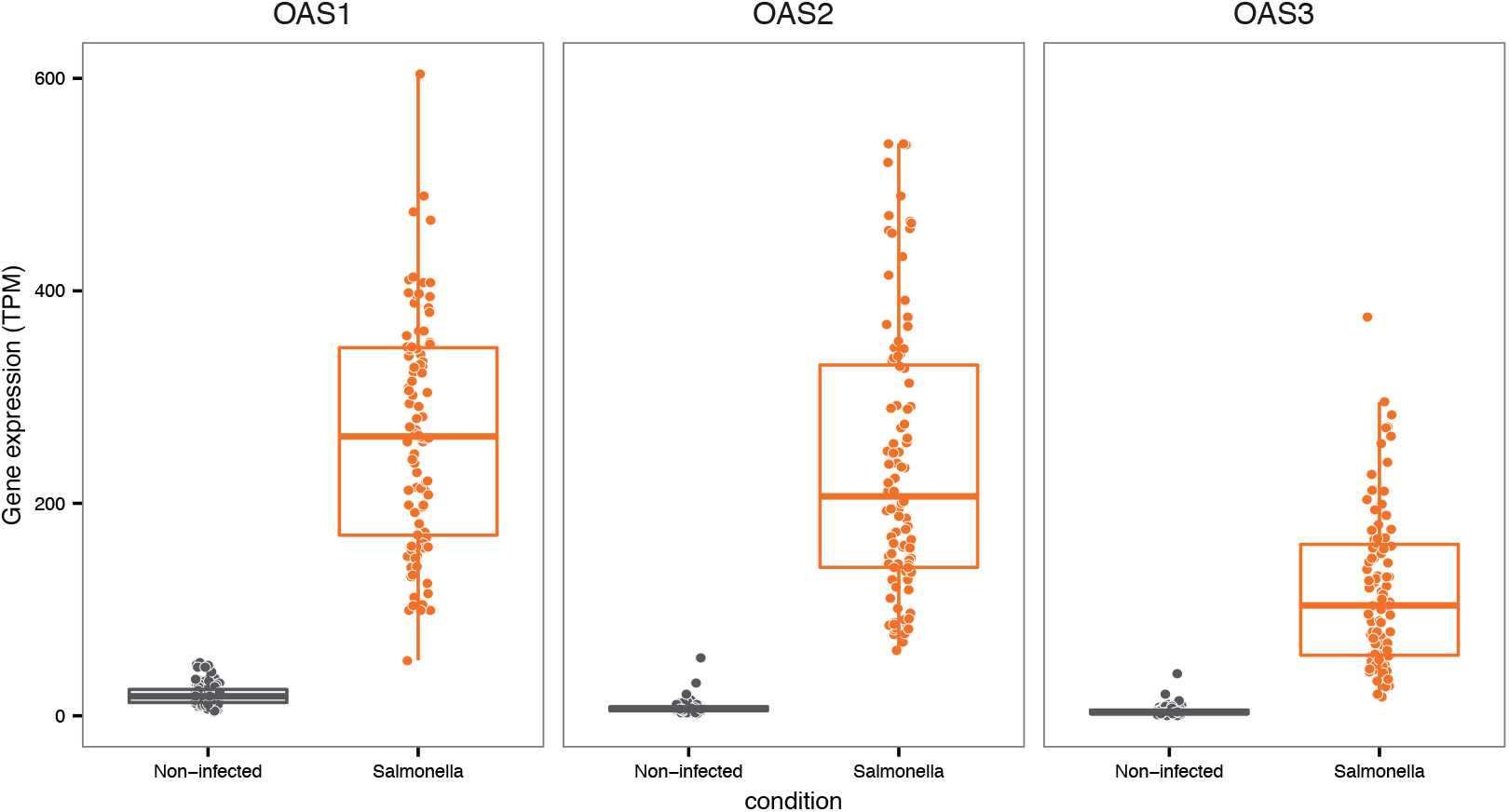
Expression levels of OAS genes in primary macrophages (European) before and after infection with Salmonella.

**Figure S6.**
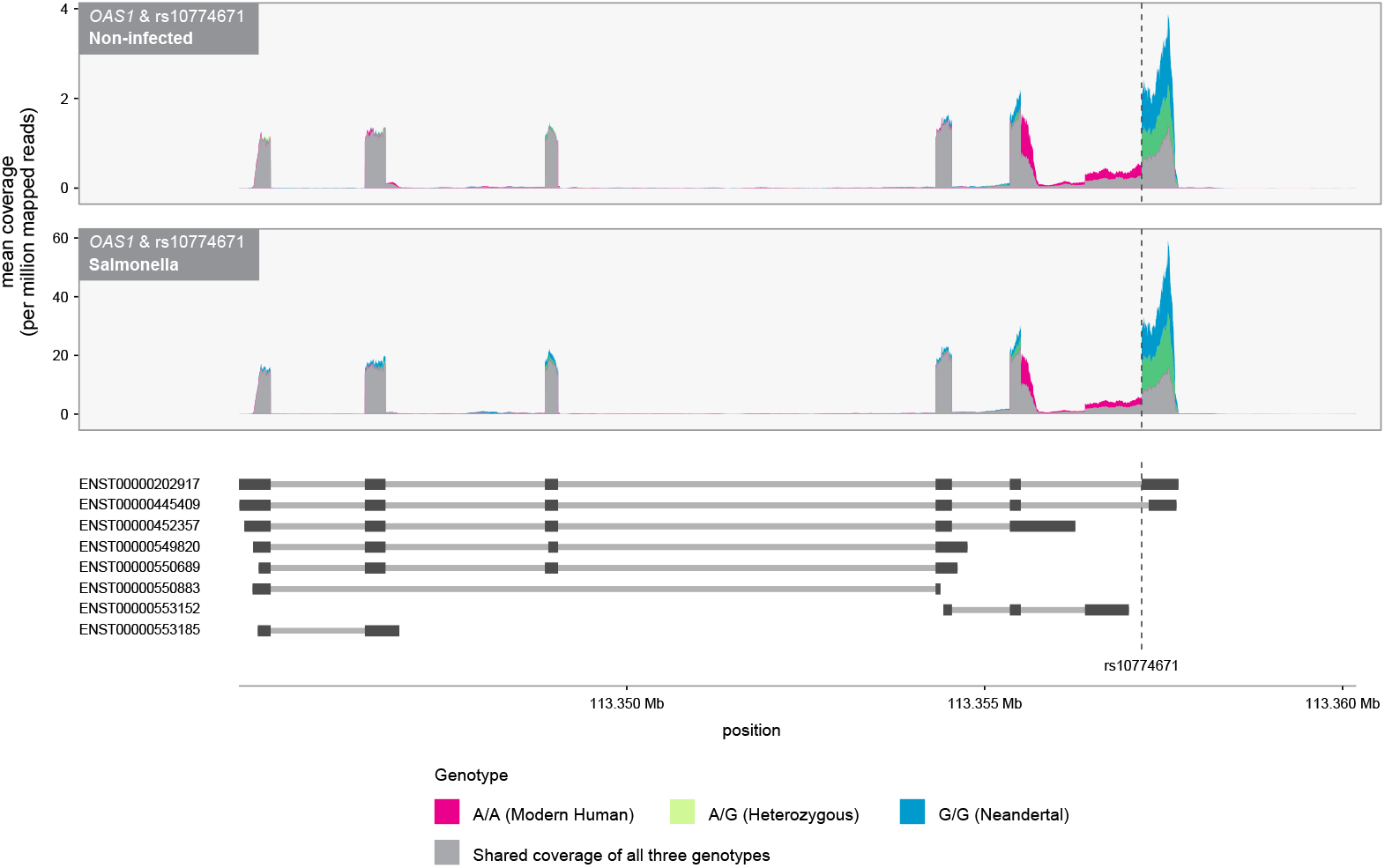
The splicing variant rs10774671 is a strong asQTL for *OAS1.* Plotted is the normalized average coverage at which each base was sequenced along the genomic regions encoding the gene *OAS1*. Individuals were stratified according to their genotype at rs10774671. Below the figure are gene models from the Ensembl database. Individuals carrying the G allele at rs10774671 (i.e. the Neandertal allele) primarily express the transcript ENST00000202917 (referred to as p46 in the text) whereas individuals carrying the A derived allele lose the splice site, which leads to the usage of a distinct isoform.

**Figure S7.**
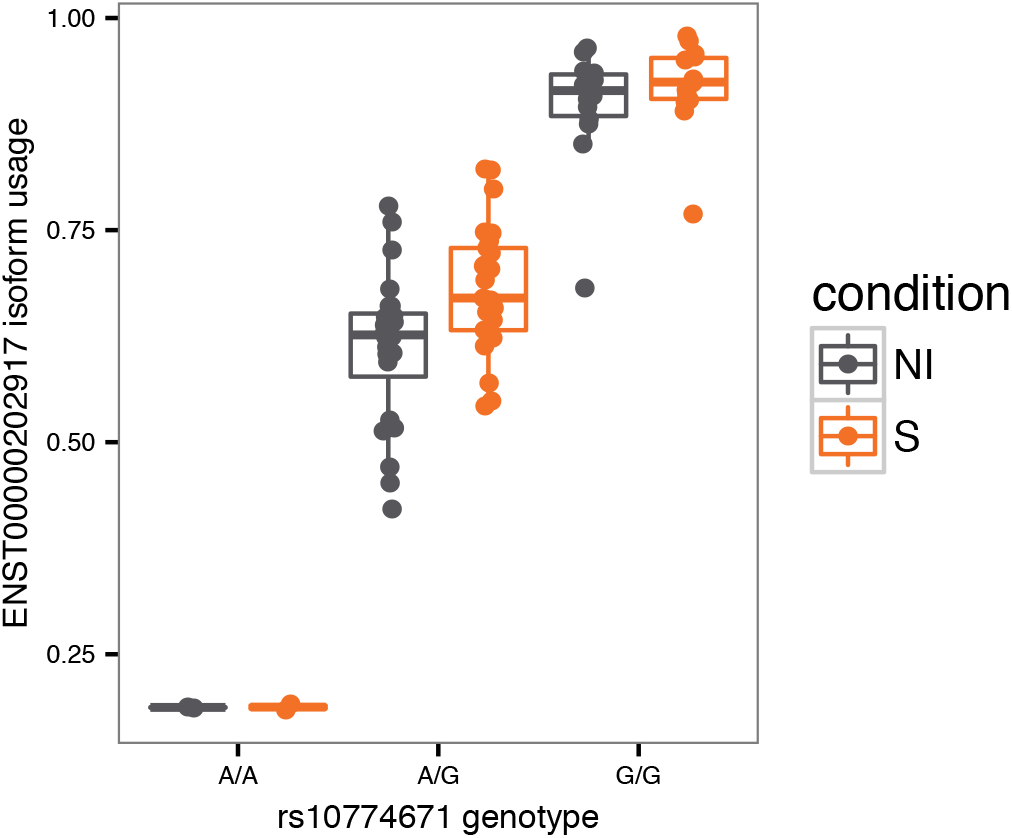
Isoform usage for transcript ENST00000202917 stratified by genotype at rs10774671 in cells derived from individuals with African ancestry. The derived (Neandertal) allele leads to increased expression of ENST00000202917 in both non-infected and *Salmonella* infected conditions.

**Figure S8.**
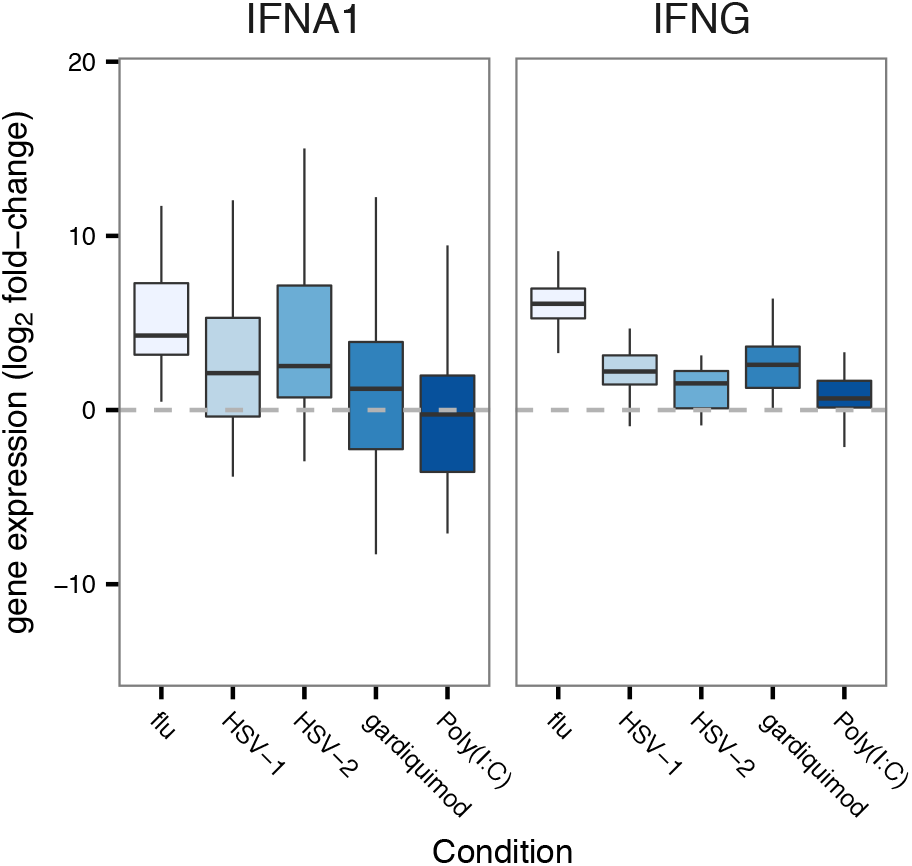
Log 2 fold induction (y-axis) of IFNA1 (type 1 interferon) and IFNG (type II interferon) in PBMCs upon stimulation with several viral agents.

**Figure S9.**
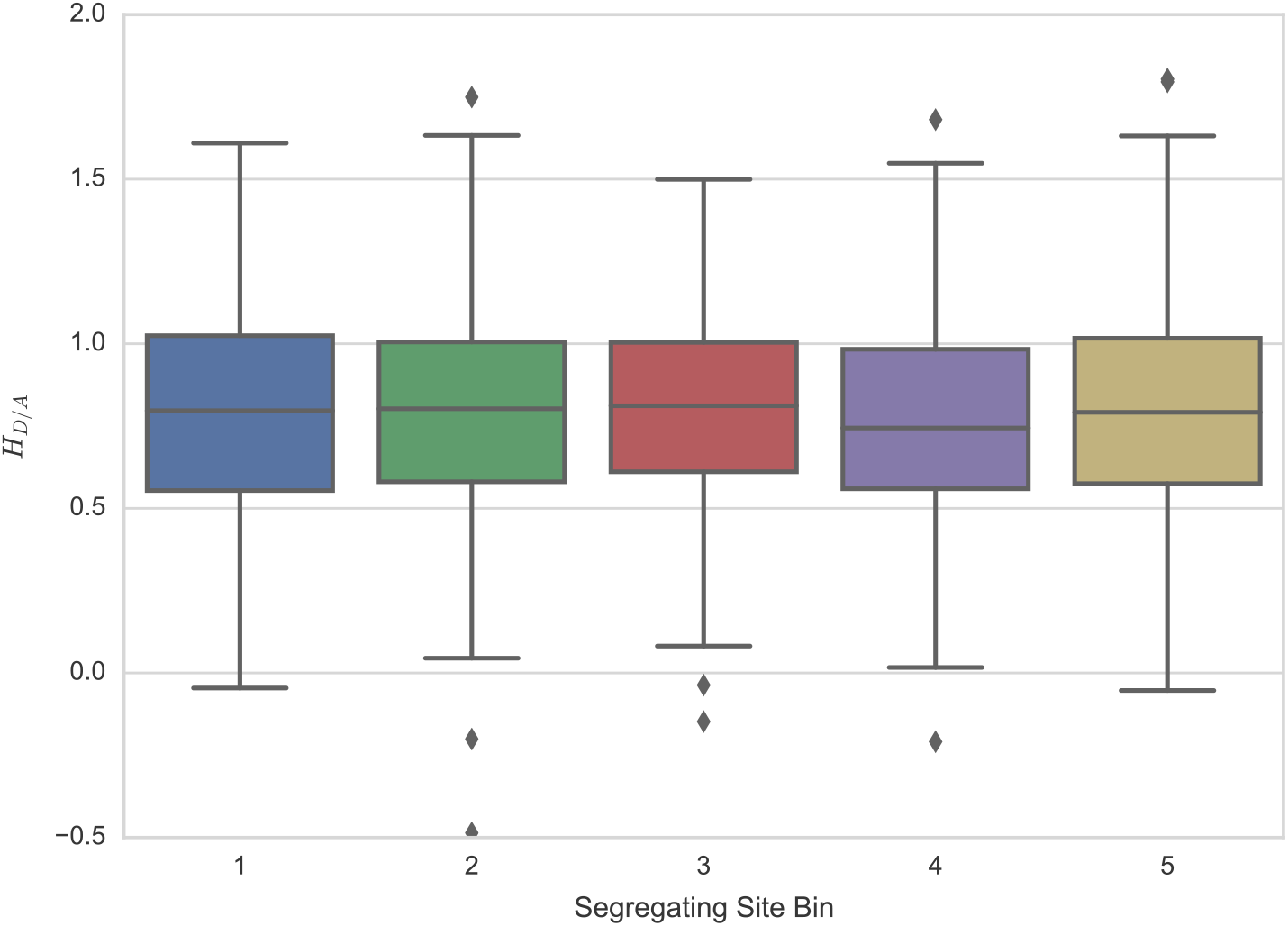
*H*_*D/A*_ plotted against quintiles of segregating sites in 1,000 simulations. The *H*_*D/A*_ statistic does vary with number of segregating sites across simulated data.

**Figure S10.**
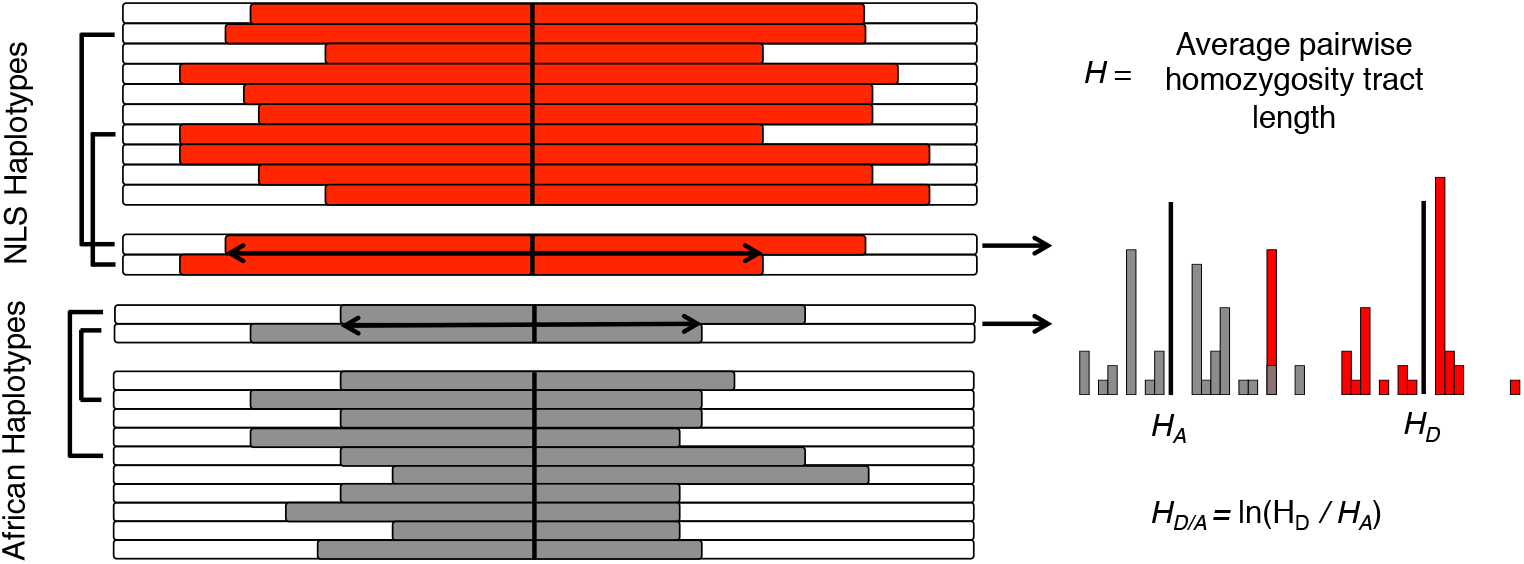
Schematic illustration of the *H*_*D/A*_ statistic. At each NLS, haplotypes are divided into two groups, based on whether they carry the Neandertal (derived) or African (ancestral) state. Within each haplotype subset, haplotype homozygosity is calculated across all pairs of haplotypes and then averaged *(H),* resulting in two values; *H*_*A*_ and *H*_*D*_ (A-ancestral, D-derived). Finally, *H*_*D/A*_ is calculated as the natural log of the ratio of derived to ancestral *H* values. Excessively high values of this statistic reflect particularly long haplotypes carrying Neandertal-derived alleles.

**Figure S11.**
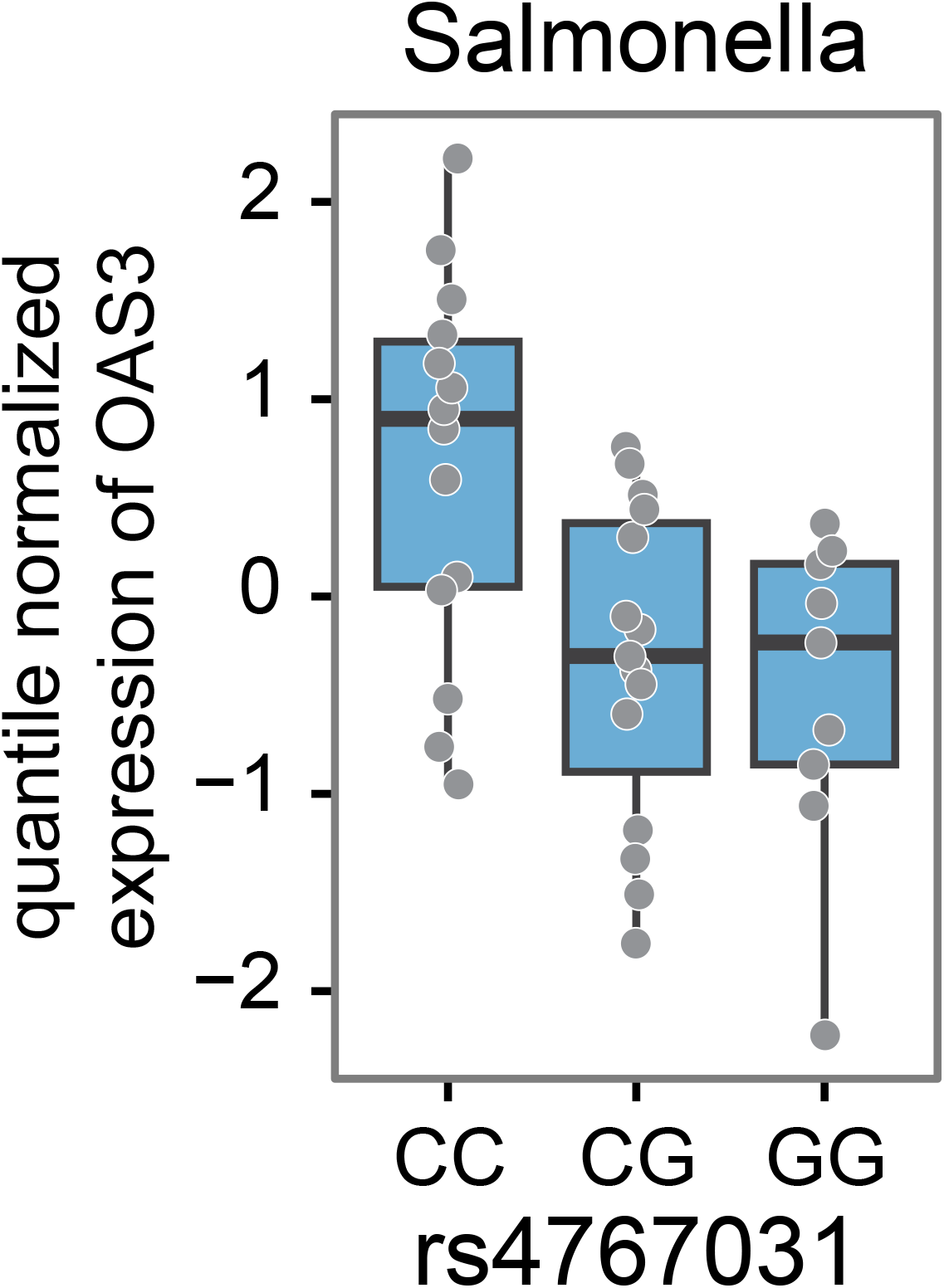
Boxplot for the association between genotypes at the NLS rs4767031 (x-axis) and the expression levels of *OAS3* in *Salmonella-infected* macrophages using real-time PCR.

